# Subspecies and sexual craniofacial size and shape variations in Japanese macaques (*Macaca fuscata*)

**DOI:** 10.1101/467456

**Authors:** Wataru Yano, Naoko Egi, Tomo Takano, Naomichi Ogihara

## Abstract

In order to investigate craniofacial size and three-dimensional shape variations independently in the Japanese macaque (*Macaca fuscata*) we used a geometric morphometries technique. A total of 55 specimens were CT scanned to generate a three-dimensional model of each cranium, and 57 landmarks were digitized to analyze the craniofacial shape variation in the Japanese macaque. The results showed that four intra-specific groups, consisting of two subspecies and the two sexes, differed in both size and shape space. In size, the cranium of the *Macaca fuscata yakui* (MFY) was smaller than that of *Macaca fuscata fuscata* (MFF) in both sexes, and female crania were smaller than male crania in both subspecies. Shape sexual dimorphisms in both subspecies were detected in the first axis of principal component analysis and were related to a relatively broad orbit, smaller neurocranium, enlarged snout, and broader temporal fossa in males. The shape differences between subspecies showed different features than those between sexes. Male subspecies shape differences were detected in the first and third axes, while those for females were in the first and second axes. Subspecies shape differences common to both sexes were a narrower orbit, relatively small neurocranium, longer snout, and postorbital constriction in MFY. Male MFY was specifically characterized by a more anterior and superior direction of snout protrusion. In contrast, female MFY showed an inferior direction of snout protrusion. Female MFY also had a taller orbit. With regard to the relationship between size and shape differences, shape sexual dimorphism for each subspecies was positively associated with size difference, but there was no such association between subspecies in either sex. Size does not seem to play an important role in subspeciation of *Macaca fuscata.*

## Introduction

The Japanese macaque (*Macaca fuscata*) ranges in the northern-most area in extant non-human primates, and inhabits the Japanese archipelago. Kuroda (1940) firstly distinguished the population in Yakushima Island from other populations based on its dwarfed body size and diagnostic pelage color and proposed the subspecies status of *Macaca fuscata yakui* (MFY) as distinct from other populations of *Macaca fuscata fuscata* (MFF). According to Nozawa et al. (1977), gene flow from the mainland of Japanese archipelago to Yakushima Island after the Marine Isotope Stages (MIS) 6–8 (0.13–0.30 Ma) should have been very rare or unlikely. Evidence from mtDNA data demonstrated that female monkeys did not migrate to Yakushima Island after 178 kya (Hayaishi and Kawamoto, 2006).

Cranial form differences between MFY and MFF have been diagnosed by using linear distance based morphometrics (craniometry: Ikeda and Watanabe, 1966; Mouri and Nishimura 2002; somatometry: Iwamoto, 1971, Hamada et al., 1996). According to those studies, regarding size variation, MFY crania are generally smaller than MFF crania. Regarding shape, Ikeda and Watanabe (1966) also pointed out that MFY has expanded zygoma, postorbital constriction, higher lamda and inion, narrower orbit, and protrusive snout, based on index comparison. Iwamoto (1971) also noted a significantly larger relative head modulus in MFY, possibly due to higher lamda, and lower cephalic index in MFY, possibly due to posteriorly positioned glabella in MFY. These conventional osteometric studies, however, leave some unanswered questions about defining statistically independent size and shape metrics. To more rigorously explore subspecies shape variations that are mathematically independent from size, we employed landmark-based three dimensional geometric morphometrics (GM). GM enables independent definitions of size and shape, and graphical presentation of the results because it preserves geometric relationships among landmarks. (Zelditch et al., 2004; Slice, 2005). The usefulness of this method has been tested in inter- and intraspecific primates size and shape analyses (e.g. O’Higgins and Dryden 1993; Frost et al., 2003; Pan et al., 2003; Schaefer et al., 2004; Cardini et al., 2008a,b). The aim of our study was to test whether or not four intraspecific groups of Japanese macaque consisting of two subspecies and the two sexes differ in their cranial morphology in size and shape.

## Methods

A total of 55 adult dried crania of Japanese macaque (14 Male MFFs, 13 Female MFFs, 14 Male MFYs, and 14 Female MFYs) were obtained from the Laboratory of Physical Anthropology, Kyoto University (Kyoto, Japan), Primate Research Institute, Kyoto University (Inuyama, Japan), and Japan Monkey Centre (Inuyama, Japan: JMC). We used adult specimens whose upper third molars were fully erupted. We did not consider the differences between wild and captive specimens for any of the four groups. MFF specimens were taken from various sites in the mainland of the Japanese archipelago. The origins of MFY specimens were from the wild Yakushima Island population and the captive population in JMC.

Each specimen was scanned with a helical computed tomography (CT) scanner (TSX-002A/4I, Toshiba Medical Systems, Tokyo, Japan) at the Laboratory of Physical Anthropology, Kyoto University. Tube voltage and current were set at 120 kV and 100 mA. Cross-sectional images were reconstructed with a pixel size of 0.20 to 0.25 mm and slice interval of 0.20 mm. The 3D surface of the cranium was then generated using a triangular mesh model with commercial software (Analyze 6.0, Mayo Clinic, Rochester, MN, USA).

We digitized a total of 57 landmarks (Fig. 1, Table 1) on the external and internal surfaces of each cranium using commercial software (Rapidform 2004, INUS Technology, Seoul, Korea). All the crania were measured by the same person (WY). As shape variances due to left-right asymmetry were not considered here, we symmetrized the positions of all landmarks using self-symmetrization (Zollikofer and Ponce de Leon, 2002). Specifically, we created the horizontally reflected specimen for each sample and superimposed them by least-square superposition to calculate the mean of each landmark coordinate, yielding the symmetrized specimen of the original. Then we analyzed the variances in the landmark positions using the geometric morphometric software Morphologika version 2.3.1 (O’Higgins and Jones, 2006). In this method, a set of landmark coordinates of each cranium were firstly scaled by centroid size (CS). CS is the square root of the sum of squared Euclidian distances from each landmark to the mean of the configuration of landmark coordinates. Subsequently, normalized landmark coordinates for each specimen were registered using the Generalized Procrustes Analysis (GPA). Thus, the landmark configuration of each cranium was represented by a single point in Kendall’s shape space. The points were projected onto a linear tangent space for subsequent statistical analyses. Principal Component Analysis (PCA) was conducted to extract principal components (PCs) of shape variations among crania (O’Higgins and Jones, 1998). Within this shape space, the relative positions of the means of the four intraspecific groups were compared. Using the software, the shape differences along the principal axes could also be visualized using 3D deformation of the wireframe connecting the landmarks.

**Fig. 1.**
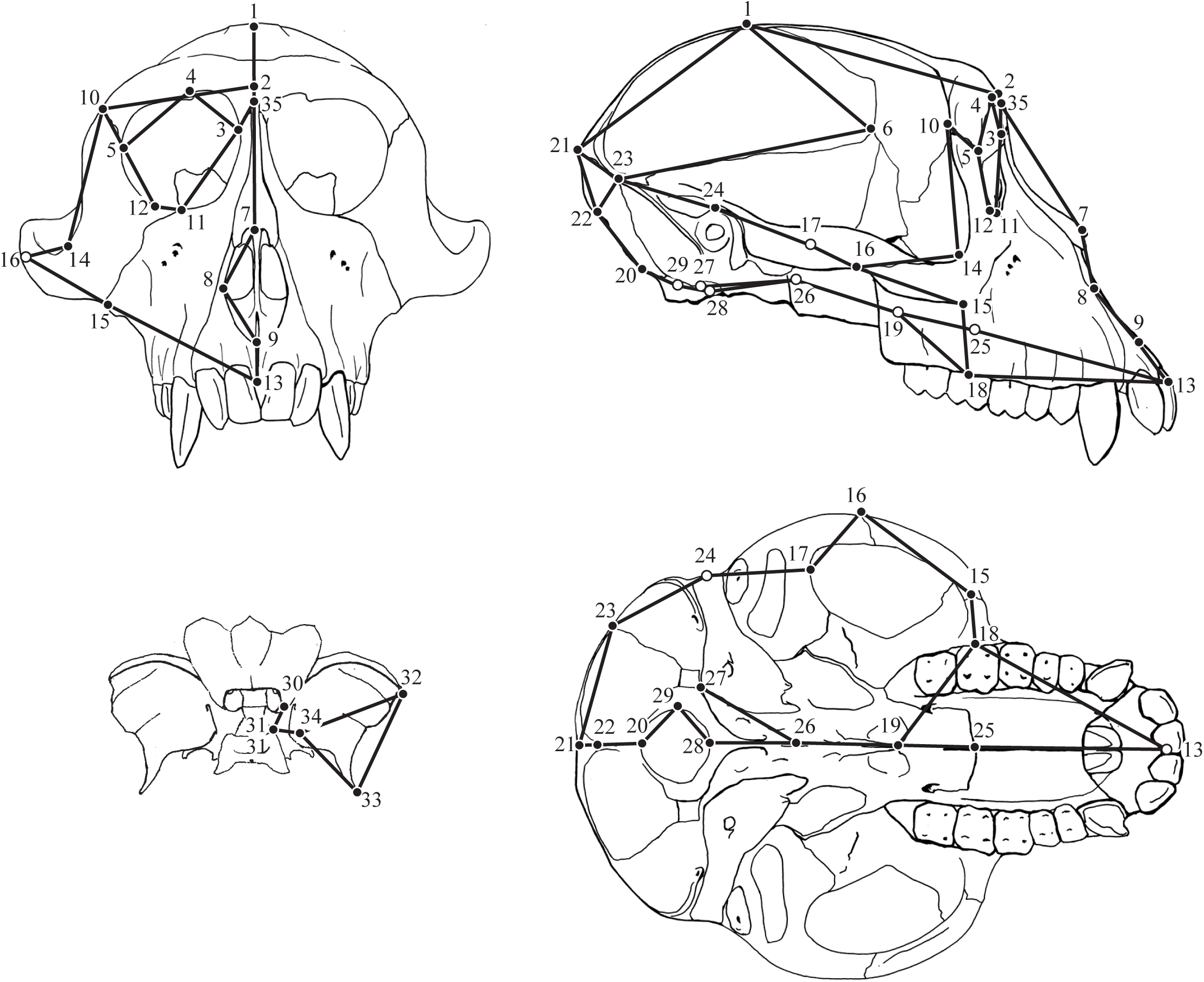
Landmarks and wireframe used in the present study. (A) Anterior view. (B) Lateral view. (C) Posterior view of sphenoid. (D) Inferior view.

**Table 1.** Landmarks used in this study.

| No. | Definition | Type |
| --- | --- | --- |
| 1 | Bregma | M |
| 2 | Supraorbitale | M |
| 3 | Maxillofrontale | B |
| 4 | Most superior point on the inferior orbital rim of frontal bone | B |
| 5 | Frontomalare orbitale | B |
| 6 | Sphenion | B |
| 7 | The ectocranial midline point on superior rim of anterior nasal aperture | M |
| 8 | Alare | B |
| 9 | Nasospinale | M |
| 10 | Frontomalare temporale | B |
| 11 | Zygoorbitale | B |
| 12 | Most inferolateral point on inferior orbital rim | B |
| 13 | Prosthion | M |
| 14 | Jugale | B |
| 15 | Zygomaxillare | B |
| 16 | Most posterior point in the temporal process of the zygomatic bone | B |
| 17 | The point in the depth of the angle between the zygomatic process and the squama of temporal bone | B |
| 18 | Midpoint on the buccal alveolar rim of second molar | B |
| 19 | Staphylion | M |
| 20 | Opisthion | M |
| 21 | Inion | M |
| 22 | The ectocranial midline point where inferior nuchal line crosses | M |
| 23 | Asterion | B |
| 24 | Auriculare | B |
| 25 | The cross-sectional point of median and transverse suture of palatine | M |
| 26 | Sphenobasion | M |
| 27 | The most anterior point of the hypoglossal canal of lateral part of occipital bone | B |
| 28 | Basion | M |
| 29 | Most lateral point on the lateral margin of the foramen magnum of lateral part of occipital bone | B |
| 30 | Most lateral point on the optic canal of lesser wing of sphenoid bone | B |
| 31 | Midpoint on the lateral side of the superior surface of the postsphenoid part of the body of sphenoid bone | B |
| 32 | Most superolateral point on the greater wing of sphenoid bone | B |
| 33 | Most infero-lateral point on the greater wing of sphenoid bone | B |
| 34 | Most inferior point on the foramen rotundum of sphenoid bone | B |
| 35 | Nasion | M |

M = midsagittal B = bilateral

The deviation from the normal distribution of CS and PC scores for each group was tested with the Shapiro Wilk test. If the normality test was passed Analysis of variance (ANOVA) tests and post-hoc Tukey’s HSD tests were employed to test for significant differences in CS and the PC scores among the four intraspecific groups. These statistical tests were performed using Statistica 2000 version 5.5 (StatSoft, Tulsa, OK, USA).

## Results

The proportions of eigenvalues of PCA are listed in Table 2. Approximately 80% of the variance was incorporated in the first fifteen PCs. Significance differences among the four intraspecific groups were found for PC1, PC2, PC3 with the ANOVA test. Other axes were not explored in this study. Proportion of eigenvalue of PC1 accounted for 20.0% of the total variance, PC2 for 11.7%., and PC3 for 9.36%. The normality test for each scores (CS, PC1, PC2, PC3) for each groups was passed demonstrating no significant deviation from normal distribution.

**Table 2.** Proportion of eigenvalues of principal compoments

| Axis | Proportion(%) | Cumulative(%) |
| --- | --- | --- |
| PC1 | 20.00 | 20.00 |
| PC2 | 11.70 | 31.70 |
| PC3 | 9.36 | 41.10 |
| PC4 | 6.17 | 47.30 |
| PC5 | 5.54 | 52.80 |
| PC6 | 5.04 | 57.80 |
| PC7 | 3.78 | 61.60 |
| PC8 | 3.42 | 65.00 |
| PC9 | 2.87 | 67.90 |
| PC10 | 2.79 | 70.70 |
| PC11 | 2.43 | 73.10 |
| PC12 | 2.05 | 75.20 |
| PC13 | 1.95 | 77.10 |
| PC14 | 1.86 | 79.00 |
| PC15 | 1.72 | 80.70 |

The four intraspecific groups clearly differed from one another in size variation represented by CS (Fig.2). The ANOVA test showed that mean scores of CS for the four intraspecific groups were not homogeneous (p<0.001), and subsequent post hoc Tukey’s HSD test revealed that males had larger CS than female for both subspecies (p<0.001), and likewise, MFF were larger than MFY for both sexes (p<0.01) (Fig.3A).

**Fig.2.**
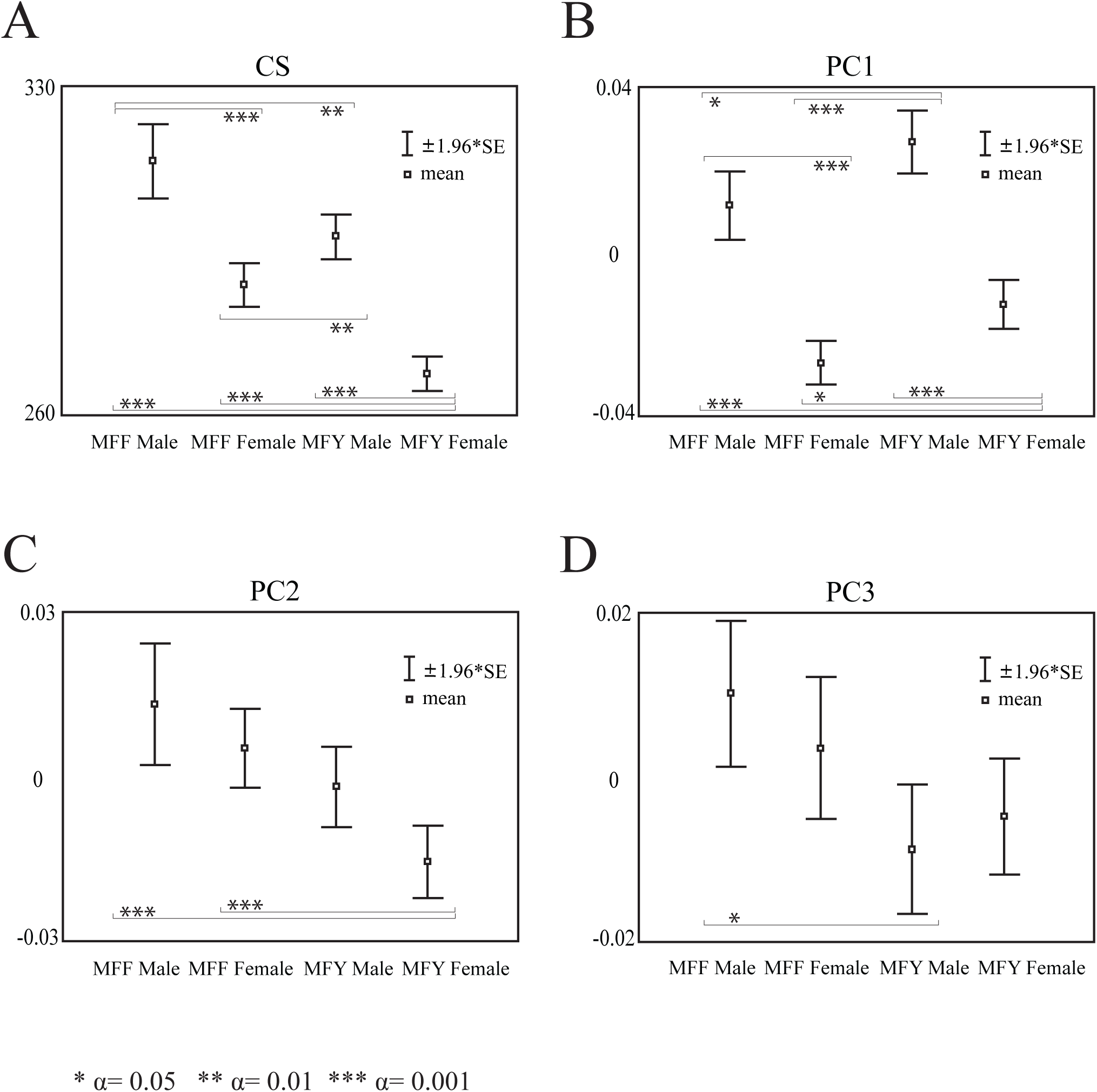
Comparison of mean scores among the four intraspecific groups. (A) Centroid size (B) PC1 (C) PC2 (D) PC3. Lines represent 95% confidence intervals.

**Fig.3.**
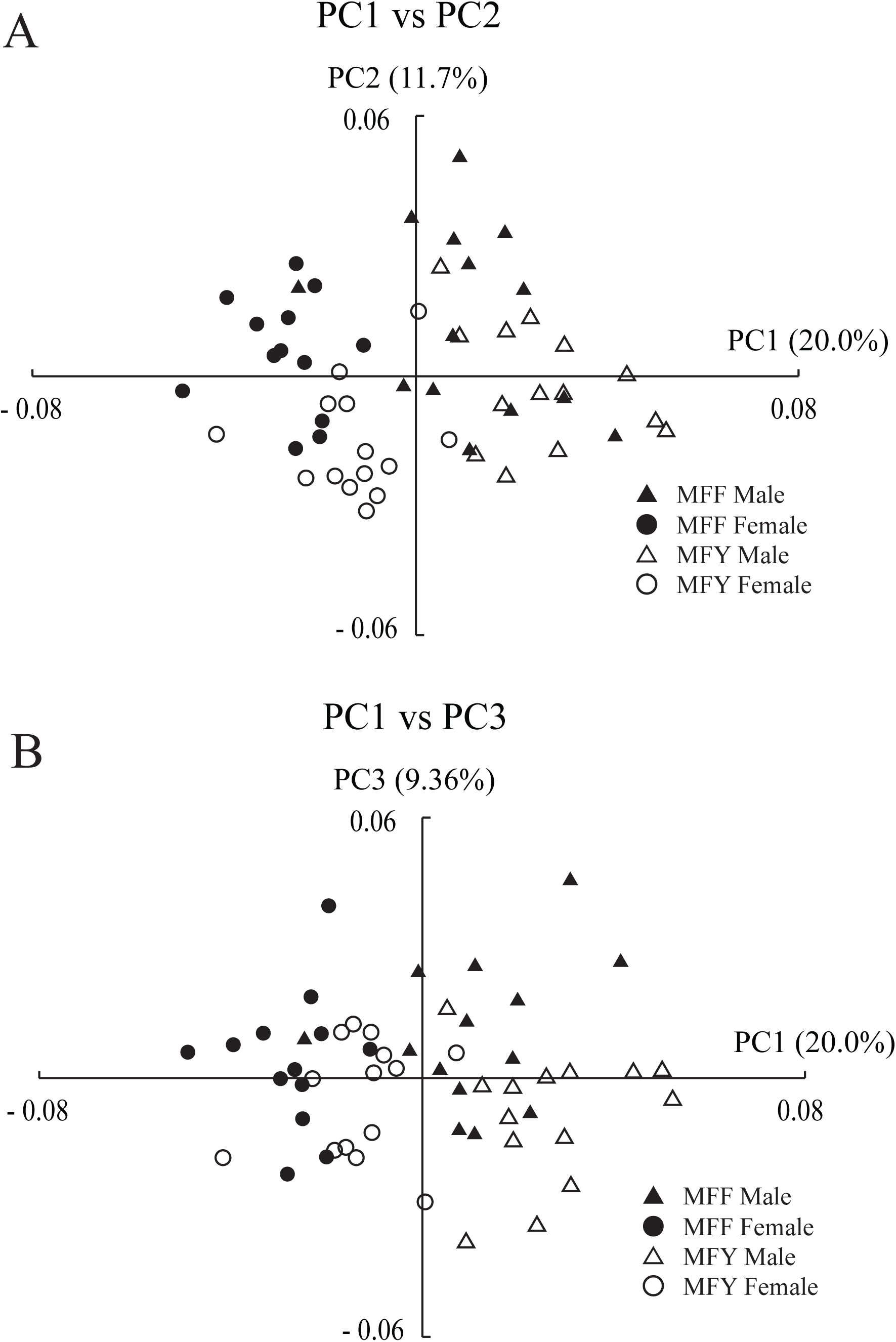
Result of principal component analysis. (A) PC1 vs. PC2. (B) PC1 vs. PC3

The shape variations are illustrated as scatter plots of PC1 versus PC2, and PC1 versus PC3, in Figure 4. Along the PC1 axis, males clearly had higher scores than females for both subspecies. Likewise, MFY had slightly higher PC1 scores than MFF for both sexes. The ANOVA test demonstrated that mean PC1 scores for the four groups were not the same (p<0.001). Tukey’s HSD test revealed that males had higher scores than females for both subspecies (p<0.001), and MFY had higher scores than MFF for both sexes (p<0.05) (Fig.3B). Besides, ANOVA also showed that mean scores for both PC2 and PC3 were not equal among the four groups (p<0.001). Tukey’s HSD test indicated that the PC2 score for MFY females was lower than that for MFF males and MFF females (p<0.01) and the PC3 score for MFY males was lower than that for MFF males (p<0.05) (Fig.3C, D).

**Fig.4.**
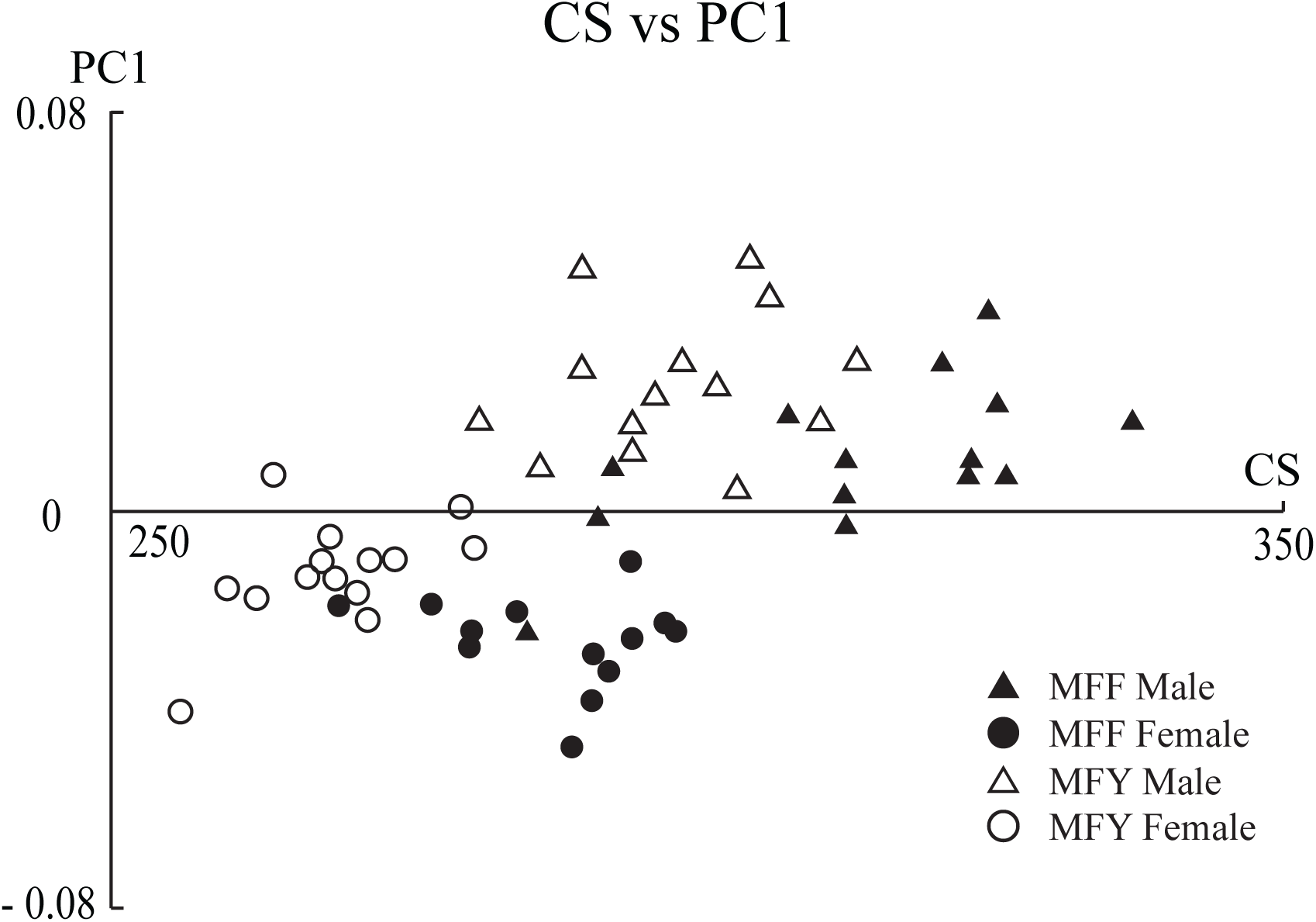
Relationships of size (CS) and PC1.

Figures 5-7 are visual presentations of 3D shape variations among the four intraspecific groups made by warping along the significant PC axes. The cranial shape was represented by the wireframe connecting landmarks. Since sexual dimorphism in shape was detected only in PC1, 3D shape variation is represented by warping the female mean (PC1= -0.02: dashed line) to the male mean (PC1=0.02: solid line) (Fig.5). The male cranium, having positive PC1, is characterized by a lower neurocranium, supero-anteriorly positioned as well as vertically tilted nuchal crest, medio-laterally expanded zygomatic arch, relatively small orbit, infero-anteriorly protruded muzzle, and relatively developed face compared to the neurocranium as viewed globally.

**Fig.5.**
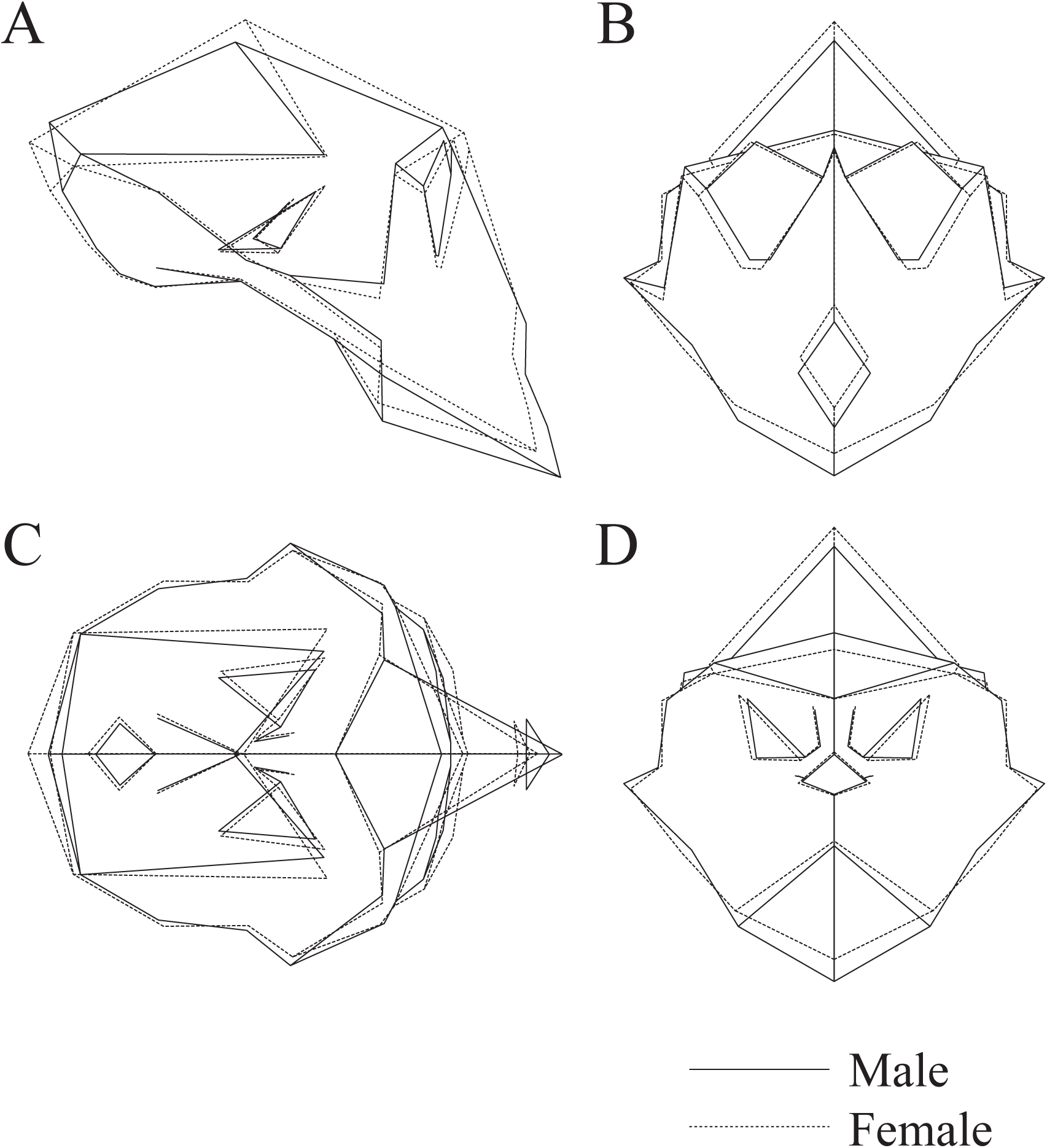
Sexual dimorphic shape difference in MFY. Solid lines, Male (PC1=0.02); Dashed lines, Female (PC1= -0.02). Shape variations are visualized with 3D deformation of the wireframe connecting landmarks

Male subspecies shape variation was visualized by warping between the mean score coordinates of MFF males (PC1=0.01, PC3=0.01: solid line) from those of MFY males (PC1=0.02, PC3= -0.01: dashed line), since male subspecies variation was along the diagonal between PC1 and PC3 (Fig.6). Likewise, subspecies shape variation in females was visualized by warping mean MFF female scores (PC1= -0.03, PC2= 0.01: solid line) from the MFY female scores (PC1= -0.01, PC2= -0.02: dashed line) (Fig.7). In both sexes, MFY tended to have a lower neurocranium, relatively vertically tilted nuchal crest, stronger post-orbital constriction, relatively narrow orbit, and relatively developed muzzle compared to the calvarium as viewed globally. One subspecies shape differences, that varied between males and females was the direction of muzzle development; the development was toward the supero-anterior direction in MFY males, while it was toward the inferior direction in MFY females.

**Fig.6.**
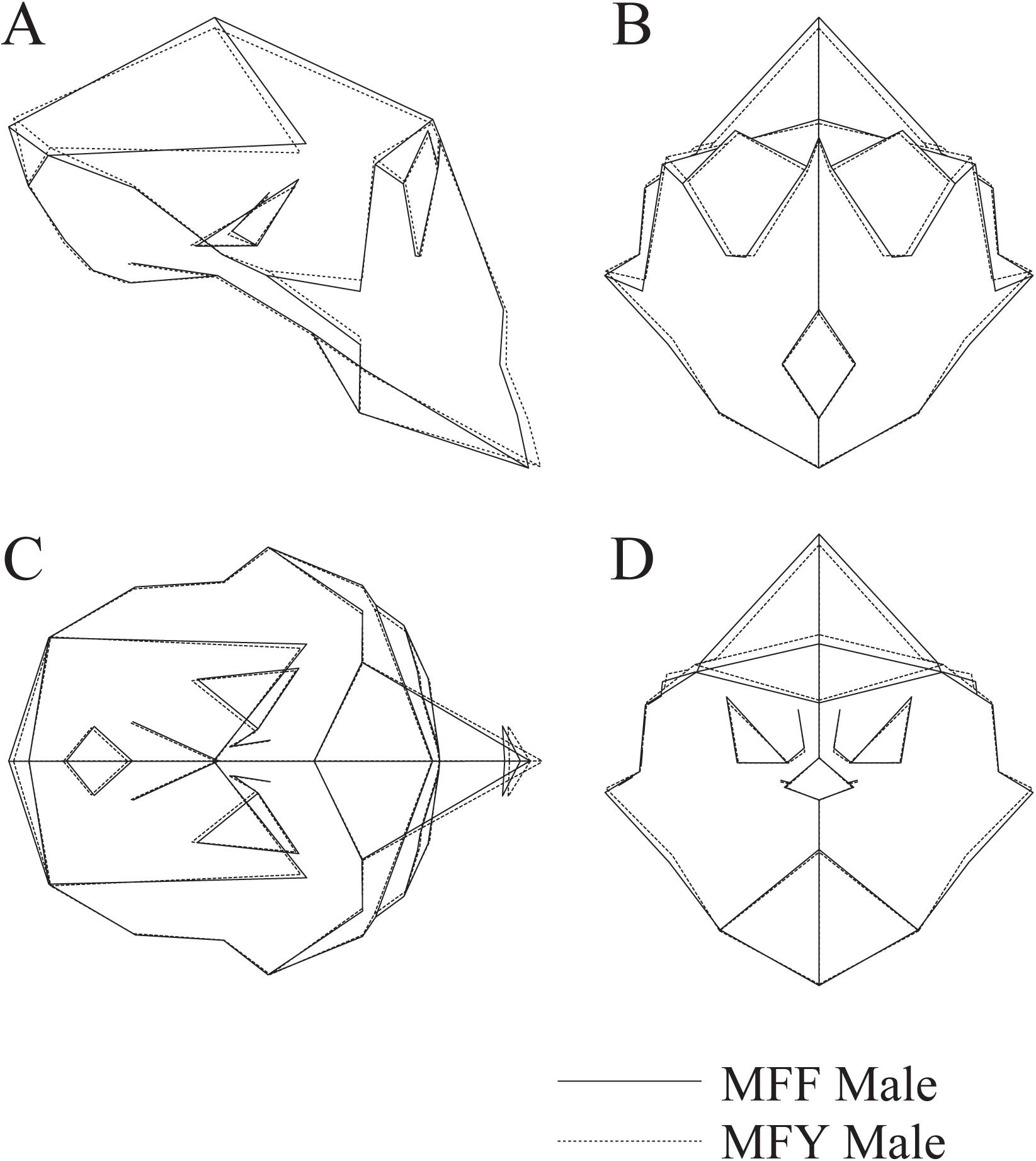
Subspecies shape differences in males. Solid lines, MFF males (PC1=0.01, PC3=0.01); Dashed lines, MFY males (PC1 = 0.02, PC3=-0.01).

**Fig.7.**
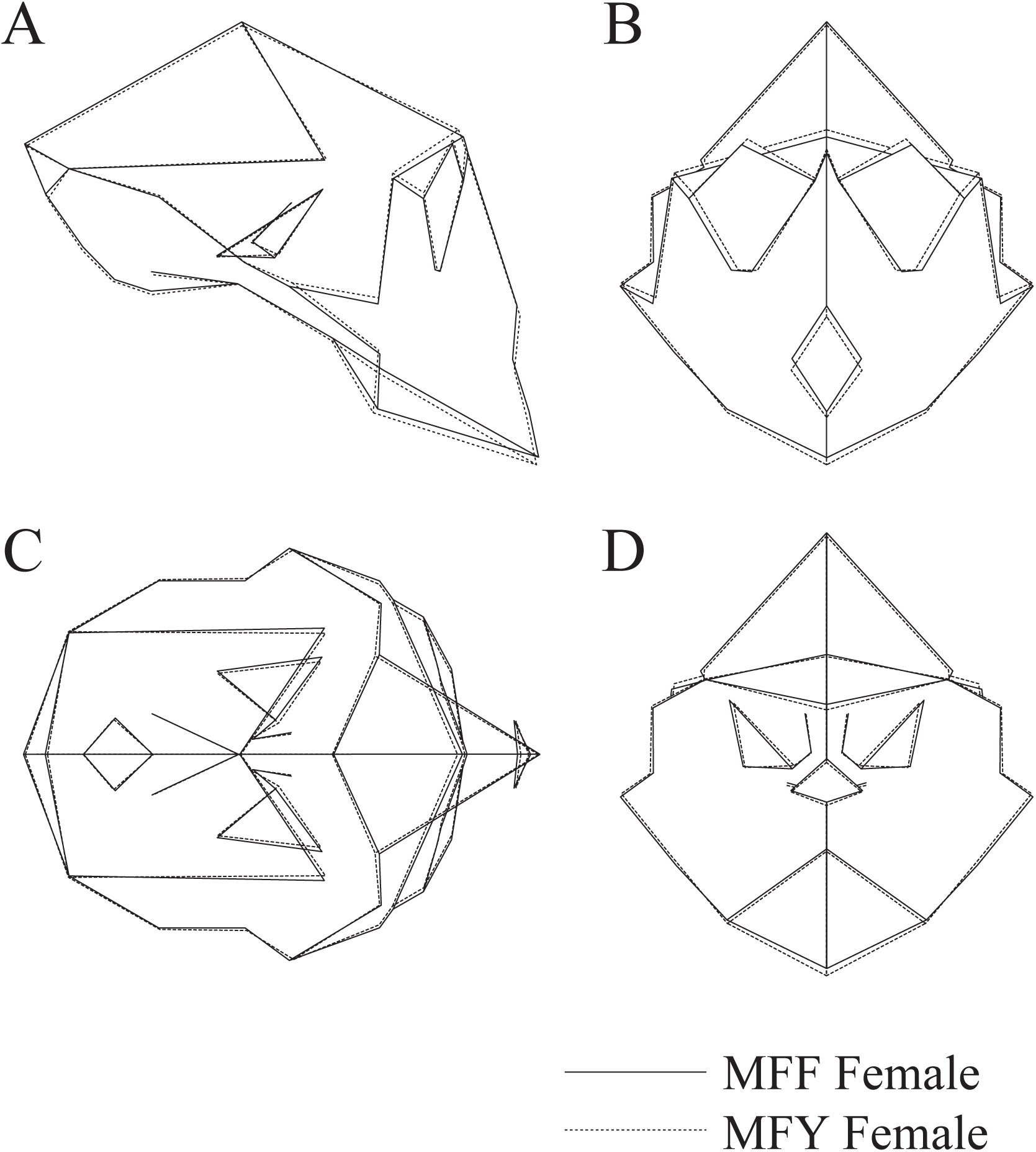
Subspecies shape differences in females. Solid lines, MFF females (PC1 = -0.03, PC2=0.01); Dashed lines, MFY females (PC1= -0.01, PC2= -0.02).

## Discussion

This study revealed that distinctive cranial size and three-dimensional shape variations exist both between the two subspecies and between the sexes of *Macaca fuscata.* The significant sexual (female<male) and subspecies (MFY<MFF) size differences we found are consistent with previous studies (Ikeda and Watanabe, 1966; Iwamoto, 1971; Hamada et al., 1996; Mouri and Nishimura 2002). The smaller cranium of MFY may be understood partly from the perspective of the insular effect (island rule). However, we cannot distinguish that effect from the effect of Bergman’s rule along latitude, which has been confirmed in this specie (Hamada et al., 1996; Kuroda 1984). Differences in size might occur via ontogenetic processes involving modification of growth rate and/or growth duration. In size sexual dimorphism, these heterochronic process appear to play a key role, as Mouri (1994) found that the adolescent growth spurt ends later in males, which leads to extension of cranial the developmental period in *Macaca fuscata*. Although the contribution of developmental processes to the subspecies variation of size is still unknown, the relative head size of MFY is already smaller at birth (Hamada, 1994).

Apparent shape differences were also recognized here between subspecies and between sexes. Shape change along PC1, and thus sexual dimorphism in this study, followed the general sexual dimorphic trend in primates (e.g. Cardini and Elton, 2002b) and might possibly be related to the canine development, which is closely related to male-male competition for mating (Plavcan, 2001). Other sexually dimorphic characters such as higher and broader occipital bun (for the insertion of nuchal muscles) and broader temporal fossa (accommodating temporal muscle passage) are related to the development of these muscles attached to the cranium. This shape sexual dimorphism is consistent with findings of a previous study obtained by using traditional morphometrics (Ikeda and Watanabe, 1966). The sexual shape variation in each subspecies appears to be correlated with size, so it is possible that shape variation can be largely explained by size difference i.e. ontogenetic scaling. To explore whether shape sexual dimorphism in the Japanese macaque is explained by ontogenetic scaling or not, however, ontogenetic data to compare ontogenetic trajectories of males and females would be necessary (e.g. Cobb and O’Higgins, 2007).

Among subspecies shape differences, the relatively vertically tilted nuchal crest, post-orbital constriction, and narrower orbit demonstrated in this study are the same as the differences reported by Ikeda and Watanabe (1966). The additional findings of the lower neurocranium and relatively developed muzzle in MFY found in this study had not been detected in any previous studies. This is because previous morphometric studies did not control for size effects and in reality compared subspecies form (size + shape) differences so that the larger muzzle in the dwarfed MFY might have been offset. It is also of interest that many parts of the characteristic traits of MFY, such as more protruding muzzle, relatively small neurocranium, postorbital constriction, and more vertically tilted nuchal plane are similar to features of the sexual dimorphism contributing to positive PC1 score for MFY in both sexes, resulting in the cranial shape in MFY appearing more “developed”, although MFY are smaller in size for both sexes. Kuroda (2002) also pointed out in his nonmetric study with adult crania that MFY is more hyperostetic than MFF. Yano et al. (2010) indicated that divergence of
ontogenetic trajectories occurs at a very early stage of fetal life, and neither pre- nor postnatal ontogenetic scaling nor heterochony alone can explain the generation of the subspecies shape differences. The early divergence of ontogenetic trajectories has also been suggested in human and non-human primates (Richtsmeier et al. 1993; Ponce de León and Zollikofer 2001; Ackermann and Krovitz 2002; Cobb and O’Higgins 2004). Although the actual ontogenetic patterns such as ontogenetic scaling and heterochrony involved in generating these subspecies differences cannot be extrapolated from the adult data in this study, shape variation is clearly not associated with size variation and size does not seem to play an important role in subspecies shape differences (Fig.4). To explore precisely when and how primates cranial shape differences are formed along ontogenetic process, comparative samples between closely related extant primates from fetus to adult will be needed.

## Acknowledgements

We wish to sincerely thank Kazumichi Katayama, Masato Nakatsukasa, Toshisada Nishida, and Daisuke Shimizu for their continuous guidance and support throughout the course of the present study. This study was supported by Japan Society for the Promotion of Science (JSPS) Grant-in-Aid for Scientific Research (B) 19370101 to NO and in part by the Global Center of Excellence Program A06 “Formation of a Strategic Base for Biodiversity and Evolutionary Research: from Genome to Ecosystem” of the Ministry of Education, Culture, Sports and Technology (MEXT), Japan.

## Reference

Ackermann RR, Krovitz GE (2002) Common patterns of facial ontogeny in the hominid lineage. Anat Rec 269:142–147

Cobb SN, O’Higgins P (2004) Hominins do not share a common postnatal facial ontogenetic shape trajectory. J Exp Zool B Mol Dev Evol 302B:302–321

Cardini A, Elton S (2008a) Variation in guenon skulls (I): Species divergence, ecological and genetic differences. J Hum Evol 54:615-37

Cardini A, Elton S (2008b) Variation in guenon skulls (II): Sexual dimorphism. J Hum Evol 54:638-47

Cobb SN, O’Higgins P (2007) The ontogeny of sexual dimorphism in the facial skeleton of the African apes. J Hum Evol 53:176-190

Foster, J B (1964) Evolution of mammals on islands. Nature 202:234–235

Frost SR, Marcus LF, Bookstein FL, Reddy DP, Delson E (2003) Cranial allometry, phylogeography, and systematics of large-bodied papionins (primates: Cercopithecinae) inferred from geometric morphometric analysis of landmark data. Anat Rec A 275A:1048-72

Hamada Y (1994) Standard Growth patterns and variations in growth patterns of the Japanese Monkeys (*Macaca fuscata*) based on an analysis by the spline function method. Anthropol. Sci. 102 (Suppl.):57-76

Hamada Y, Watanabe T, Iwamoto M (1996) Morphological variations among local populations of Japanese Macaque (*Macaca fuscata*). In Shotake T and Wada K (eds) Variations in the Asian Macaques. Tokai University Press, Tokyo pp 97-115

Hayaishi S, Kawamoto Y (2006) Low genetic diversity and biased distribution of mitochondrial DNA haplotypes in the Japanese macaque (*Macaca fuscata yakui*) on Yakushima Island. Primates 47:158–164

Ikeda J, Watanabe T (1966) Morphological studies of *Macaca fuscata* III Craniometry. Primates 7:271-288

Iwamoto M (1971) Morphological studies of *Macaca fuscata* IV Somatometry. Primates 12:151-174

Kuroda N (1940) A monograph of the Japanese mammals exclusive of sirenia and cetacean. Sanseido Co, Tokyo, pp 311 (in Japanese)

Kuroda S (1984) Morphological characters and evolution of Yakushima macaques. Monkey 198/199:14-17 (in Japanese)

Kuroda S (2002) Microevolution of cranial bone morphology of Macaca fuscata in the postglacial period Non-metrological mutant characters of cranial bone of *Macaca fuscata*. Asian Paleoprimatology 2:115-125 (in Japanese)

Lomolino, M V (1985) Body size of mammals on islands: the island rule re-examined. American Naturalist 125:310–316

Mouri T, Nishimura T (2002) Craniometry of Adult Male Japanese Macaques from the Yakushima, Koshima and Kinkazan Islands. Primate Research 18(1):43-47 (in Japanese, English summary)

Nozawa K, Shotake T, Minezawa M, Kawamoto Y, Hayasaka K, Kawamoto S, (1991) Population genetics of Japanese monkeys: III Ancestry and differentiation of local populations. Primates 32:411-435

O’Higgins P, Dryden I (1993) Sexual dimorphism in hominoids: further studies of craniofacial shape differences in Pan, Gorilla, and Pongo. J Hum Evol 24: 183-205

O’Higgins P, Jones N (1998) Facial growth in Cercocebus torquatus: an application of three dimensional geometric morphometric techniques to the study of morphological variation. J Anat 193:251-272

O’Higgins P, Jones N (2006) Tools for statistical shape analysis. Hull York Medical School available from http://wwwyorkacuk/re/fme/resources/software.htm

Pan R, Wei F, Li M (2003) Craniofacial variation of the Chinese macaques explored with Morphologika. J Morph 256:342-348

Plavcan (2001) Sexual Dimorphism in Primate Evolution. Yearbook of Am J Phys Anthropol 44:25–53

Ponce de León MS, Zollikofer CPE (2001) Neanderthal cranial ontogeny and its implications for late hominid diversity. Nature 412:534–538

Richtsmeier JT, Corner BD, Grausz HM, Cheverud JM, Danahey SE (1993) The role of postnatal growth pattern in the production of facial morphology. Syst Biol 42:307–330

Schaefer K, Mitteroecker P, Gunz P, Bernhard M, Bookstein F (2004) Craniofacial sexual dimorphism patterns and allometry among extant hominids. Ann Anat 186:471-478

Slice D (2005) Modern morphometrics in physical anthropology. Kluwer Acad Plenum, New York

Yano W, Egi N, Takano T, Ogihara N 2010 Prenatal ontogeny of subspecie variation in the craniofacial morphology of the Japanese macaque (*Macaca fuscata*). Primates 51:263-271

Zelditch M, Swiderski D, Sheets H, Fink W (2004) Geometric morphometrics for biologists: a primer. Elsevier, San Diego

Zollikofer CPE, Ponce de León MS (2002) Visualizing patterns of craniofacial shape variation in Homo sapiens. Proc R Soc Lond B 269:801-807

